# Improved representation of sequence Bloom trees

**DOI:** 10.1101/501452

**Authors:** Robert S. Harris, Paul Medvedev

## Abstract

**Motivation:** Algorithmic solutions to index and search biological databases are a fundamental part of bioinformatics, providing underlying components to many end-user tools. Inexpensive next generation sequencing has filled publicly available databases such as the Sequence Read Archive beyond the capacity of traditional indexing methods. Recently, the Sequence Bloom Tree (SBT) and its derivatives were proposed as a way to efficiently index such data for queries about transcript presence.

**Results:** We build on the SBT framework to construct the HowDe-SBT data structure, which uses a novel partitioning of information to reduce the construction and query time as well as the size of the index. Compared to previous SBT methods, on real RNA-seq data, HowDe-SBT can construct the index in less than 36% of the time, and with 39% less space, and can answer small-batch queries at least five times faster. We also develop a theoretical framework in which we can analyze and bound the space and query performance of HowDe-SBT compared to other SBT methods.

**Availability and implementation:** HowDe-SBT is available as a free open source program on https://github.com/medvedevgroup/HowDeSBT.

**Contact:** Paul Medvedev

**Supplementary information:** Supplementary text and figures available as single Supplementary file.

## 1 Introduction

Public read databases such as the Sequence Read Archive (SRA) contain a treasure trove of biological information and have the potential to become a ubiquitous community resource by enabling broad exploratory analyses. For example, given a long nucleotide sequence, which experiments in the database contain reads matching it? More concretely, which human RNA-seq experiments from the SRA contain a transcript of interest? Unfortunately, there does not exist a way for today’s biologist to answer such a question in a reasonable amount of time. Tapping into the potential of these databases is hampered by scalability challenges and will require novel approaches from the algorithm community.

The computational problem falls into the widely-studied category of string alignment problems (Gusfield, 1997; Mäkinen *et al*., 2015). However, it differs in several regards. The strings to be matched are rarely present in the database in their entirety. Instead, sequencers produce many highly fragmented copies of the desired string, each subjected to potential sequencing error. Furthermore, the scale of the databases makes traditional sequence alignment methods, such as SRA-BLAST (Camacho *et al*., 2009), inadequate (Solomon and Kingsford, 2016).

The seminal paper of Solomon and Kingsford (2016) demonstrated that the above transcript question can be simplified to a question of approximate *k*-mer membership. Each experiment can be viewed as a collection of its constituent *k*-mers, and the biological question can be answered by finding all experiments which contain a high percentage of the *k*-mers in the query transcript. They demonstrated that this approach, even if only done approximately, is a good proxy for the answer to the transcript question. It also lends itself to answering a more broad range of questions, such as SNP presence, viral contamination, or gene fusion (Bradley *et al*., 2017). Their work has opened the door to a slew of data structures implementing various *k*-mer indices, roughly falling into two categories.

The first category of approaches are based on the Bloofi (Crainiceanu and Lemire, 2015) data structure. Each experiment’s *k*-mers are first stored in a Bloom filter (Bloom, 1970), an efficient but lossy data structure for storing sets. The experiments are then grouped into a hierarchical structure (i.e. tree) based on their similarity, where each leaf corresponds to the *k*-mers in an experiment. Each internal node represents the union of the *k*-mers in its descendants. The tree allows an efficient search for experiments matching a given *k*-mer profile by pruning non-promising branches. Bloofi was first adapted to the sequencing context by the Sequence Bloom Tree (SBT) data structure (Solomon and Kingsford, 2016), and further work improved the representation of the internal nodes (Sun *et al*., 2018; Solomon and Kingsford, 2017) and the clustering of the tree topology (Sun *et al*., 2018).

The SBT approaches aggregate *k*-mer information at the level of an experiment. The second category of approaches aggregate experiment information at the level of the *k*-mers. In such an approach, each query *k*-mer is independently looked up in an index to retrieve information about which experiments contain the *k*-mer (Holley *et al*., 2015; Muggli *et al*., 2017; Almodaresi *et al*., 2017; Mustafa *et al*., 2018; Pandey *et al*., 2018; Yu *et al*., 2018; Bradley *et al*., 2017; Almodaresi *et al*., 2018; Holley and Melsted, 2019). In this context, experiments are referred to as colors, and such a data structure is sometimes called a colored de Bruijn graph. These approaches are complementary to the SBT and the best choice depends on the particular properties of the queries and the dataset, such as the sharedness of *k*-mers between experiments.

In this paper, we make two main contributions. First, we develop an alternative way to partition and organize the data in an SBT such that it becomes more compressible and faster to query. To demonstrate the performance advantages of our method, called HowDe-SBT, we compare it on real data to the previous SBT methods. We also propose and explore a culling procedure to remove non-informative nodes from the tree and create a non-binary forest (see Supplementary).

Second, we introduce a theoretical framework which allows us to prove bounds on the performance of HowDe-SBT in comparison with Split-SBT (abbreviated as SSBT)(Solomon and Kingsford, 2017). Previous papers in the field have focused on experimental metrics for comparison, but, while these are very valuable and necessary, they can vary greatly depending on the dataset or the system used. Theoretical bounds can deepen our understanding of why algorithms perform well and can drive the development of better methods. In this paper, we derive an information theoretic bound on the space used by an SBT (Theorem 1) and quantify the number of bit lookups necessary for a query (Theorem 2).

## 2 Preliminaries

Let *x* and *y* be two bitvectors of the same length. A bitvector can be viewed as a set of the positions that are set to 1, and in this view, the set union (intersection, respectively) of *x* and *y* is equivalent to bitwise OR (AND, respectively). We write the bitwise AND between *x* and *y* as *x* ∩ *y*, and the bitwise OR as *x ∪ y*. The bitwise NOT operation on *x* is written as 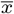. The set difference of *x* and *y* is written as *x \ y* and can be defined as 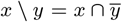. The empty set is represented as a bitvector of zeros. The universal set, denoted by *ξ*, is represented as a bitvector of ones. Given that the fraction of 1s in *x* is *p*, the *empirical entropy* of *x* is defined as *H*(*p*) = -(*p* log_2_ *p* + (1 – *p*) log_2_(1 – *p*)).

A *Bloom filter* (BF) is a bitvector of length b, together with q hash functions, *h*_1_,…, *h_q_*, where *b* and *q* are parameters. Each hash function maps a *k*-mer to an integer between 0 and *b* — 1. To add a *k*-mer *x* to the set, we set the position *h_i_* (*x*) to 1, for all *i*. To check if a *k*-mer *x* is in the set, we check that the position *h_i_*(*x*) is 1, for all *i*. Note that a false positive may occur, i.e. *x* may have never been added but all its corresponding positions were still set to 1 and it is considered to be contained in the BF. In this paper, we restrict the number of hash functions to be *q* = 1 (as is done in other SBT approaches).

Next, consider a rooted binary tree *T*. The parent of a non-root node *u* is denoted as *parerat(u)*, and the set of all the leaves of the subtree rooted at a node *u* is denoted by *leaves(u)*. Let *children(u)* refer to the child nodes of a non-leaf node *u*. Define *aracestors(u)* as 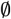 if node *u* is the root, and *parerat(u)* ∪ *aracestors*(*parerat(u)*) otherwise. That is, *aracestors(u)* are all the nodes on the path from *u* to the root, except *u* itself.

Suppose that there is a Bloom filter associated with each leaf of *T*. Then, define *B*_∪_ (*u*) for a leaf node *u* as its associated BF and *B*_∪_ (*u*) for an internal node as 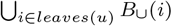. Note that *B*_∪_ (*u*) of an internal node *u* can be equivalently defined as 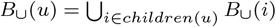. Define the intersection of leaf BFs in the subtree rooted at a node *u* as 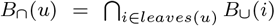. Equivalently, at internal nodes, 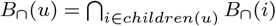.

In this paper, we solve the following problem:

### Database input

A database *D* = {*D*_1_,…, *D_n_*}, where each *D_i_* is a BF of size *b* that represents experiment *i*. Typically, *D_i_* contains all the non-erroneous *k*-mers that appear in the reads of experiment *i*.

### Query input

A multi-set of *k*-mers *Q* (called the query), and a threshold 0 < *θ* ≤ 1.

### Query output

The set of experiments whose *D_i_* contains at least a fraction *θ of* the query *k*-mers, i.e. {*i*: |{*x* ∈ *Q*: *x* exists in *D_i_*}| ≥ *θ* · |*Q*|}

Note that in this formulation, we assume that D (along with parameter b) is already given to us. How to choose b and construct D from raw reads was already described in Solomon and Kingsford (2016).

## 3 Representation and querying

Initially, we determine a tree topology T using the clustering algorithm of Sun *et al.* (2018) as a black-box. A *topology* is a binary tree with a bijection between its leaves and the experiment BFs in our database; a topology does not yet have bitvectors assigned to the internal nodes. We now show how to assign these bitvectors — *B*_det_ and *B*_how_— and how they can be used to answer a query.

Conceptually, each node *u* represents the set of experiment BFs corresponding to *leaves(u)*. We observe that some positions are *determined* in *u*: they have the same value in each of *leaves(u)*. Moreover, for the positions that are determined, we know exactly *how* they are determined. Formally, we define

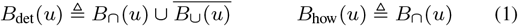

Note that Bhow is intended to only be informative for those positions that are in *B*_det_. Figure 1 shows an example of a tree and its *B*_det_ and *B*_how_ representation.

**Fig. 1.**
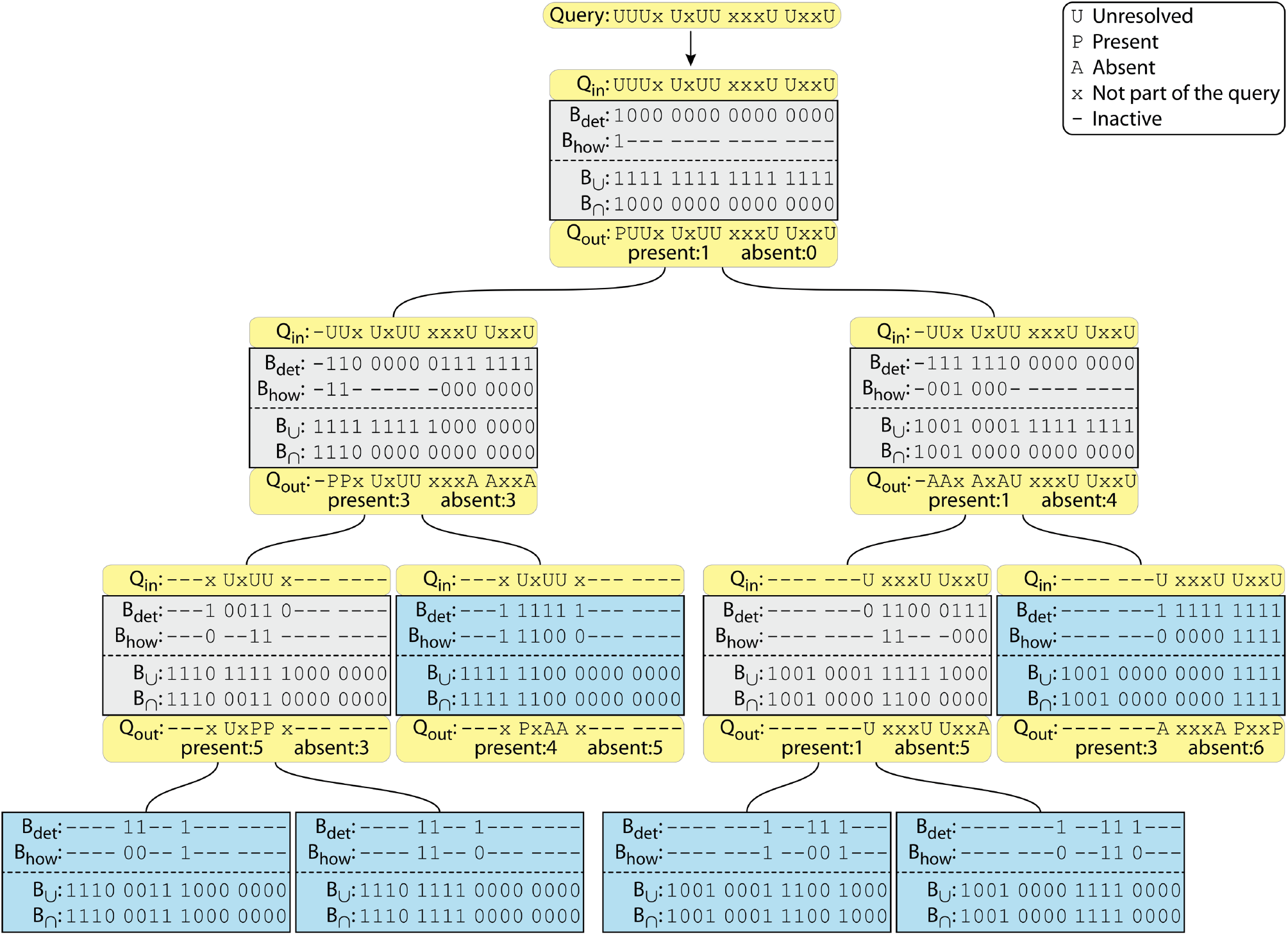
Example HowDe-SBT on *D* = {1110 0011 1000 0000; 1110 1111 0000 0000; 1111 1100 0000 0000; 1001 0001 1100 1000; 1001 0000 1111 0000; 1001 0000 0000 1111}. Each box represents a node of the tree, with leaves shown in blue and internal nodes in gray. For conciseness, the boxes show *B*_det_, *B*_how_, *B*_∪_, and *B*_∩_, but do not explicitly show 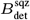 and 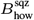. The yellow boxes demonstrate the processing of an example nine *k*-mer query with *θ* = 0:5. The initial query is shown at the top, with the bits corresponding to the query *k*-mers shown with a U. The query flows down the tree, i.e. it is processed in a preorder traversal. The query state presented to a node is *Q*_in_ and the result from the node is *Q*_out_. Some bits are resolved as either present (P) or absent (A) across the subtree. Such bits are counted in the corresponding totals and removed from further processing (hyphen) in descendants. Processing along a branch halts if either total reaches the threshold needed to determine the outcome. In this example, the search along any branch can be terminated if either the present counter or the absent counter exceeds 5. In this example, *Q* matches the first two experiments.

Having *B*_det_ and *B*_how_ at each node can enable the following efficient query search; it is essentially the same strategy as in Sun *et al.* (2018) and Solomon and Kingsford (2017), but adapted to the HowDe-SBT tree. When we receive the set of query *k*-mers *Q*, we hash each one to determine the list of BF positions corresponding to *Q* (recall that our BF uses only one hash function, hence each *k*-mer corresponds to just one position). We call this list the *unresolved* positions. We also maintain two counters: the number of positions that have been determined to be 1 *(present)*, and the number of positions determined to be 0 (*absent*). These counters are both initially 0. We then proceed in a recursive manner, starting at the root of the tree. When comparing *Q* against a node *u*, each unresolved position that is 1 in *B*_det_(*u*) is removed from the unresolved list, and the corresponding bit in *B*_how_(*u*) determines which counter, present or absent, is incremented. If the present counter is at least *θ*|*Q*|, we add *leaves(u)* to the list of matches and terminate the search of *u*’s subtree. If the absent counter exceeds (1 – θ) |*Q*|, *Q* cannot match any of the descendant leaves so we terminate the search of *u*’s subtree. If neither of these holds, we recursively pass the two counters and the list of unresolved positions down to *u*’s children. When we reach a leaf, the query unresolved list will become empty because *B*_det_ is all ones at a leaf, and the algorithm will necessarily terminate. Figure 1 shows an example of a query execution.

We observe that some bit positions will never be looked at during a search, as follows. First, if a position is determined at a node *v*, it will be removed from the query unresolved list (if it was even there) after node *v* is processed. We say that this position is inactive in *v*’s descendants, since a search will never query that position. Formally, a position is *active in B*_det_(*u*) if it is not determined at its parent (equivalently, at any of its ancestors). Second, the only positions that are queried in *B*_how_(*u*) are those that are active and set to one in *B*_det_ (*u*). We say these positions are *active in B*_how_(*u*). Formally, we can define

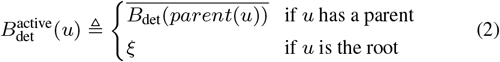

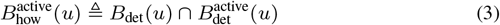

Bits that are inactive are wasteful, since they take space to store but are never queried. We remove these bits, forming (usually shorter) bitvectors comprised of only the active bits. Formally, let 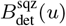 be *B*_det_(*u*) with all the inactive bits removed, and we let 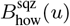 be *B*_how_(*u*) with all the inactive bits removed. 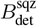 and 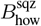 are further compressed with the general purpose RRR compression algorithm (Raman *et al*., 2007), and these compressed bitvectors are what, in the end, constitute our index. We note that since the removal of inactive bits changes the indices into the bitvectors, the query algorithm has to be modified accordingly by using rank and select. Since this is done in essentially the same way as in Solomon and Kingsford (2017), we omit the details here.

## 4 Analysis of savings compared to previous work

In this section, we show the connection of our representation to previous approaches and analyze the theoretical improvements. The structuring of our bitvectors can be viewed as an extension of the approach in Solomon and Kingsford (2017), which is called SSBT. The SSBT representation approach subsumes the representation of Sun *et al.* (2018), so we focus our comparison on SSBT. Briefly, SSBT uses the same approach of having a tree where the bitvectors at a node *u* represent, for each bit position *x*, whether *x* is 1 in all, none, or some of the *leaves(u).* It also marks a bit position as inactive if it can never be reached during a query. The SSBT bitvectors are called *u*_sim_ and *u*_rem_. The topology of the SSBT is computed differently from HowDe-SBT, and, as was shown in Sun *et al.* (2018), it results in a poorly clustered tree. However, since we want to focus on the improvements solely due to bitvector representation, we will assume that the SSBT is constructed using the same topology as HowDe-SBT. In this case, the relationship of SSBT to HowDe-SBT can be summarized as a one-to-one relationship between all possible bitvector states, shown in Figure 2.

**Fig. 2.**
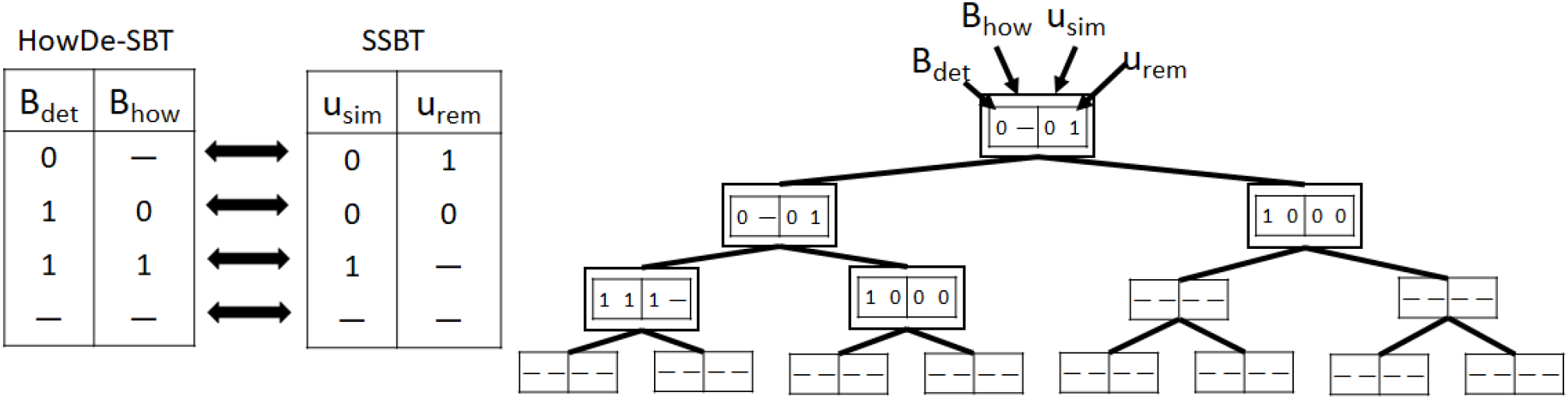
The left tables show the correspondence between the possible states of SSBT and HowDe-SBT bitvectors. A dash represents an inactive bit. The tree is an example of a bit position *x* and its values in both representations. The nodes of *T_x_* have a double border. In this example, *n_x_* = 5 and *ℓ_x_* = 3 because *T_x_* has 5 nodes and 3 leaves. Further, *s_x_* = 1 = 3 because the how bit is active in exactly three nodes (corresponding to the leaves of *T_x_*) and set to 1 in exactly one of them.

The intuition which guided our design of HowDe-SBT, relative to SSBT, was 1) minimizing the number of active positions and 1 bits (to improve space), and 2) minimizing the number of bit lookups performed during a query (to improve speed). To try to theoretically quantify this improvement, we derive a savings rate per bit position x, in terms of 1) *n_x_*, the number of nodes where *x* is active in *B*_det_, and 2) *s_x_,* the percentage of nodes where *x* is active in Bhow that have a 1.

Let *T* be the tree topology. We are not able to directly derive the savings rate for *T* after RRR compression, but we instead rely on Shannon’s information compression bound. For a bitvector *a* that is generated by *a* 0^th^ order Markov model and that has a fraction *p* of 1s, the best that a lossless compression algorithm can achieve is |*a*|*H*(*p*) bits. While in practice this bound might be beaten because our bitvectors are not generated by a 0^th^ order Markov chain, it is still a useful proxy for the compressibility (and in any case the RRR compression that we use does not compress beyond the 0^th^ order bound (Raman *et al*., 2007)). Let 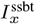 (respectively, 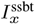) denote Shannon’s information bound for storing all the bit values at *x* in *T* using HowDe-SBT’s representation (respectively, SSBT’s representation).

### Theorem 1.

*Let* 0 ≤ *x* < *b be a bit position. Then*,

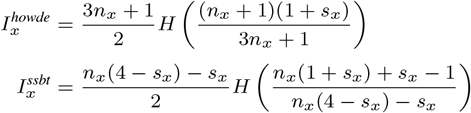

Proof. Let *T_x_* be the subtree of *T* containing all nodes where *x* is active in *B*_det_. Observe that *T_x_* is indeed a subtree and it contains exactly the nodes where *x* is determined and their ancestors. The total number of nodes in *T_x_* is *n_x_*. Let *ℓ_x_* be the number of leaves of *T_x_*. Note that since every internal node of *T* has two children, *ℓ_x_* = (*n_x_* + 1)/2, for all *x*. Note that the nodes at which the Bhow bits are active are exactly the leaves of *T_x_*, and *s_x_* is the percentage of these that are 1.

Now, the nodes for which *x* is active in *B*_det_ are exactly the nodes of *T_x_*, and thus *x* contributes *n_x_* active *B*_det_ bits. The only time *B*_how_ is active at *x* is when *B*_det_ is set to 1, which is exactly at the leaves of *T_x_*. Hence, the total number of active bits is *n_x_* + *ℓ_x_* = (3*n_x_* + 1)/2. Next, we count the number of active bits that are set to 1. The *B*_det_ bits set to one are exactly at the leaves of *T_x_*. There are *ℓ_x_* active *B*_how_ bits, of which a fraction *s_x_* are set to one. Hence, the number of active bits set to one is *ℓ_x_*(1 + *s_x_*) = (*n_x_* + 1)(1 + *s_x_*)/2.

To prove the statement about 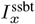, we use the equivalences in Figure 2 as a guide. The number of active positions in the *u*_sim_ vectors is *n_x_* and in the *u*_rem_ vectors is *n_x_* – *s_x_ℓ_x_*. In sum, the number of active positions is 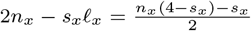. The number of active bits in *u*_sim_ that are set to 1 is *ℓ_x_s_x_* and the number of active bits in *u*_rem_ that are set to 1 is *n_x_* — *ℓ_x_*. Hence the number of 1 bits in *S* is 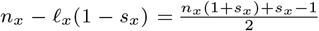.

### Theorem 2.

*Suppose both the* SSBT *and* HowDe-SBT *are built from the same tree topology T. Then the number of bit lookups necessary to resolve a bit position *x* is* (3*n_x_* + 1)/2 *in* HowDe-SBT, *and* (*n_x_*(4 — *s_x_*) — *s_x_*)/2 *in* SSBT.

Proof. Consider HowDe-SBT. For every internal node of *T_x_*, we only make one lookup to *B*_det_, resulting in *n_x_* — *ℓ_x_* lookups. At the leaves of *T_x_*, we must also look at *B*_how_, resulting in 2*ℓ_x_* lookups. The total number of lookups is then *n_x_* + *ℓ_x_* = (3*n_x_* + 1)/2. For SSBT, we must always check both *u*_sim_ and *u*_rem_ at every node, with one exception: at a leaf of *T_x_*, we do not need to check urem if usim is 1. Hence, the number of lookups is 2*n_x_* — *ℓ_x_ s_x_* = (*n_x_*(4 — *s_x_*) — *s_x_*)/2.

We can measure the percent improvement in the space bound as 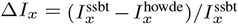, shown in Figure 3(a). In the limit, Δ*I_x_* approaches between 9% and 14% for *s_x_* ≤ 0.75. Similarly, we can measure the percentage improvement in the number of lookups as 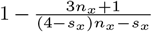 which in the limit goes to 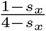. The improvement in lookups is thus between 0 (for *s_x_* = 1) and 25% (for *s_x_* = 0); for *s_x_* = 0.5, it is 14%.

In large scale applications, our theoretical analysis can be simplified by assuming that *n_x_* goes to the limit. However, HowDe-SBT can be applied in different settings, and some of these settings (like a private patient cohort) may in fact not be very large. Our detailed analysis can be used to determine at which point a dataset is large enough for the asymptotic effects to kick in; e.g. Figure 3(a) indicates that the space savings reaches a stable point roughly at = 50.

We caution, however, that our analysis does not automatically translate to total improvements when all the positions are considered jointly. The total data structure size depends a lot on the structure of the input. That is reflected by the distribution of *n_x_* and *s_x_* in the real data, whether or not they are correlated, the entropy of all the bits together, rather than separated by position, and the higher-order entropy of the bitvectors. These effects can be analyzed using real data, which we do in Section 6.

**Fig. 3.**
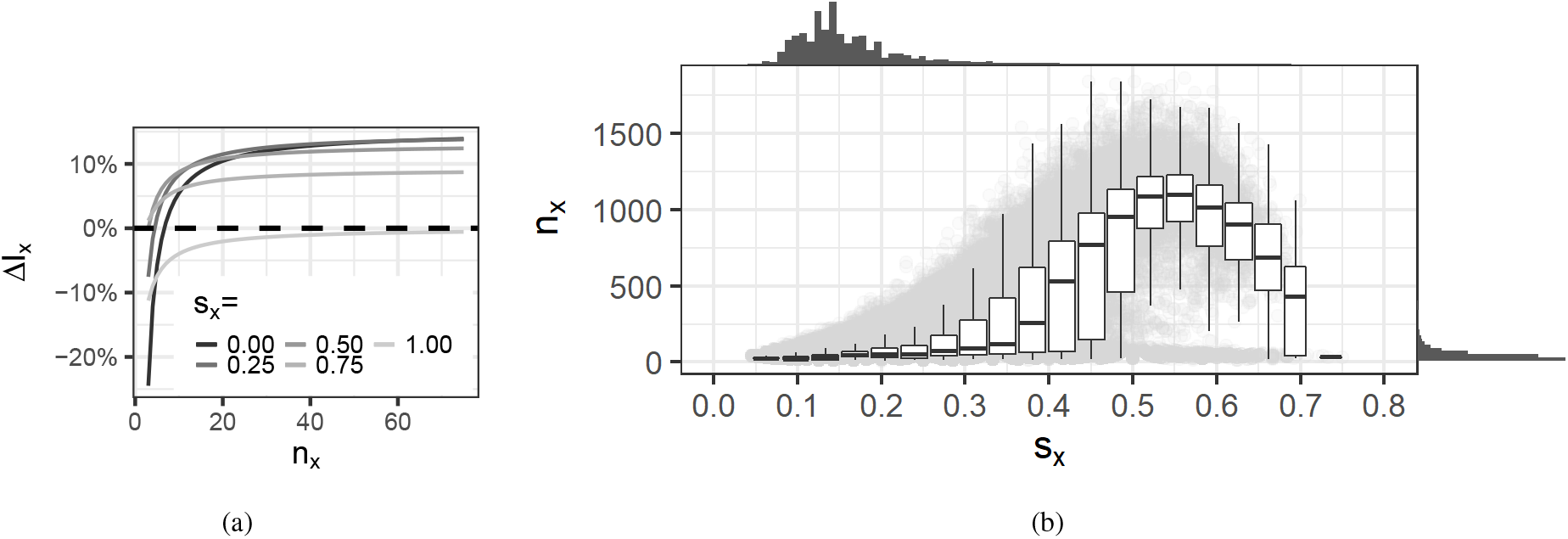
(a) Δ*I_x_* as a function of *n_x_*, for various values of *s_x_*. (b) The distributions of and dependencies between *s_x_* and *n_x_* on the real data. We use a random subsample of 1 mil bit positions for the plot, but omit those positions that are determined at the root (i.e. *n_x_* = 1). These make up 15% of all values, and they all have *s_x_* = 0. Each of the remaining positions are shown as a gray dot in the background. The *s_x_* values are binned and the distribution of *n_x_* within each bin is shown as a box plot. On the top (respectively, right) of the plot is the histogram of *s_x_* (respectively, *n_x_*).

## 5 Construction algorithm

In this section, we describe an algorithm to compute 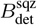 and 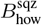 for a given topology and prove its correctness. Our algorithm makes a pass through the tree using a postorder traversal. At each node *u*, it 1) computes *B*_det_ and *B*_how_ of *u*, and 2) computes 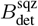 and 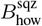 of each of its children using their *B*_det_ and *B*_how_. As the final step, 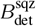 and 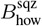 are computed for the root node.

The base case of the algorithm is, for each leaf *u*, to set *B*_det_ (*u*) ← *ξ* and *B*_how_(*u*) ← *B*_∪_(*u*). For an internal node *u*, we first construct *B*_∩_ (*u*) and *B*_∪_ (*u*) using the operations:

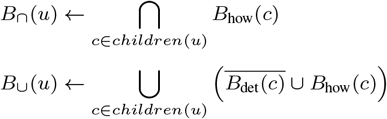

Then, *B*_det_ (*u*) and *B*_how_ (*u*) are constructed by directly applying their definitions (Equation (1)). Once *B*_det_(*u*) and *B*_how_ (*u*) have been computed, we compute the active bits in *B*_det_ (*c*) and *B*_how_ (*c*), for each child *c* of *u*:

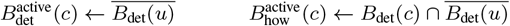

Next, we make a linear scan through *B*_det_ (*c*) and *B*_how_ (*c*) and copy over only the bits that are set in 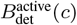 and 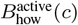, respectively, thus gyrating 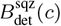 and 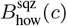.

These are finally RRR compressed and written to disk.

After the postorder traversal completes, we make the final step to compute the 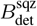 and 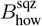, for the root *u*. It is essentially the same process as for the internal nodes, but the active bits are set as:

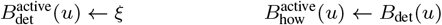

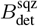 and 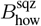 are then computed as before, removing the inactive bits during a linear scan followed by RRR compression.

We note that the 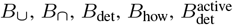, and 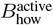 bitvectors are not stored after construction is complete. To save memory, these can be discarded right after their use; however, it is necessary to save some of the *B*_det_ and *B*_how_ vectors. Specifically, we must save *B*_det_(*u*) and *B*_how_(*u*) between the visit to *u* and the visit to *u*’s parent. During this interval they can be saved in memory or, in order to save memory, they can be written to disk and reloaded to memory as needed (the default behavior). When *u*’s parentis visited, 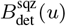 and 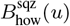 are computed and *B*_det_(*u*) and *B*_how_(*u*) are no longer needed.

The following theorem shows that our algorithm is correct and that its runtime is linear in the total size of the input bitvectors.

### Theorem 3.

*Given a database of n Bloom filters, each of size b, and a tree topology, our method constructs 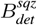 and 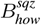 for all nodes in 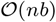 time*.

Proof. We first prove the runtime. Our bitvector construction algorithm operates on each node twice — once when the node is visited, and a second time to finish the node when its parent is visited. (The root is a special case — its finishing stage occurs at the end of the algorithm.) At each stage, it performs a constant number of bitwise operations which each takes 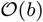 time. Similarly, the linear scan to remove inactive bits can be done in 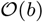 time by simply copying the active bits to a new vector.

Next, we prove correctness. We will need the following technical lemma:

### Lemma 1.

*The following properties are true*:

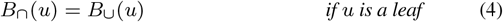

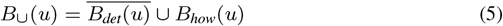

Proof. Equation (4) trivially follows from the definition of *B*_∩_. For Equation (5), first observe that *B*_∩_ (*u*) and 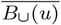 are disjoint (since 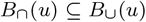). Combine this with the definition of *B*_det_(*u*), and we get 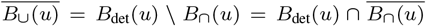. Negating both sides, we get 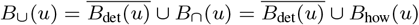

We first show the correctness of computing *B*_det_ and *B*_how_ when we visit *u*. For a leaf *u*,

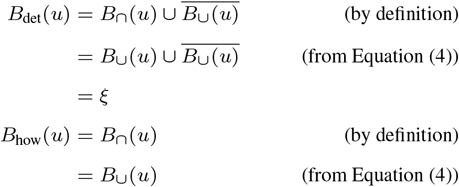

If *u* is an internal node, then it is enough to show that *B*_∩_ and *B*_∪_ are computed correctly.

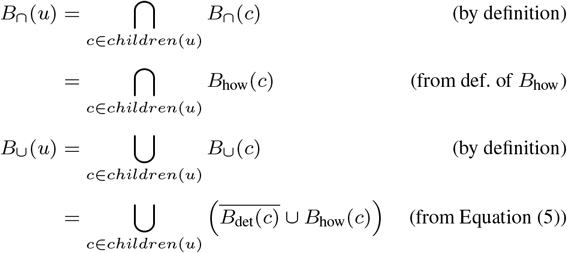

To show the correctness of computing 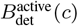 and 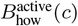, recall their definitions from Section 3. It is straightforward to see that our algorithm computes 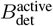 and 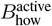 exactly according to these definitions, both for the case of children and for the case of the root. Finally, the algorithm correctly computes 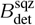 and 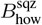 because it simply applies their definitions.

We note that the complexity of constructing the tree topology was not studied in Sun *et al.* (2018), but the obvious implementation would take Ω(*n*^2^) time. Though it takes negligible time on currently tested datasets, it may become the bottleneck in the future if n increases by several orders of magnitude.

## 6 Results

### 6.1 Experimental setup

To evaluate HowDe-SBT, we compared it against its two predecessors, AllSome-SBT (Sun *et al*., 2018) and SSBT (Solomon and Kingsford, 2017). All experiments were run on an Intel Xeon CPU with 512 GB of RAM and 64 cores (at 2.10 GHz). Our tool is open source and available for free through https://github.com/medvedevgroup/HowDeSBT. All details about how the tools were run, including parameters, together with datasets needed to reproduce our results, are available on https://github.com/medvedevgroup/HowDeSBT/tree/master/reproduce.

We used the same data for evaluation as Solomon and Kingsford (2016), except that we removed experiments that did not have reads longer than *k*. There were 66 experiments removed in this way. A 67^th^ experiment was removed because the corresponding BF in Solomon and Kingsford (2016) was empty. Together, the removed experiments had a total of 675 million reads. The resulting dataset contained 2,585 human RNA-seq runs from blood, brain, and breast tissues, compromising all relevant human RNA-seq datasets in the SRA at the time of Solomon and Kingsford (2016). For each file, we filtered out any *k*-mers that occurred less than a file-dependent threshold. We used the thresholds from Pandey *et al.* (2018), for consistency purposes. We used *k* = 20 and a Bloom filter size of *b* = 2 · 10^9^ (as in previous work), and *k*-mer counting was done using Jellyfish (Marçais and Kingsford, 2011).

To study query performance, we created four types of queries: a single transcript, a batch of ten transcripts, a batch of 100, and a batch of 1000. Transcripts were picked arbitrarily from Gencode (ver. 25) transcripts that are at least *k* nt long. We created 100, 10, 3, and 3 replicates for each type of query, respectively. We include batches of multiple transcripts in our tests because SBT performance is known to depend on batch size (Solomon and Kingsford, 2016). The idea of combining multiple queries in a batch and then processing them at each node simultaneously was first described by Solomon and Kingsford (2016) and is implemented in all SBT methods, including HowDe-SBT. We use a value of *θ* = 0.9 for all experiments. Note that because the output of HowDe-SBT is identical to SSBT and AllSome-SBT, we do not need to compare their accuracy; moreover, a comparison of SBT accuracy relative to exact methods like Mantis was also already explored in Solomon and Kingsford (2016).

### 6.2 Performance comparison

Table 1 shows the time and space taken to construct the index. The index could be created in less than 36% of the time and with 39% less space for HowDe-SBT than for all other approaches. The faster construction time of HowDe-SBT over AllSome-SBT was due to a combination of having to handle much smaller bitvectors during construction (as reflected by the smaller index size) and software engineering improvements. Otherwise, the construction algorithms of the two methods differ only by the specific bitvector operations applied during the tree traversal.

**Table 1.** File sizes (in GiB) and build times. The intermediate space refers to the initial experiment Bloom filters. The construction time is the time to build the index from the BFs and does not include *k*-mer counting. All times shown are on one processor.

|  | HowDe-SBT | AllSome-SBT | SSBT |
| --- | --- | --- | --- |
| construction time | 9h | 25h | 57h |
| intermediate space | 602 | 602 | 602 |
| index size | 14 | 142 | 23 |

It is important to note that the construction times in Table 1 do not include the pre-processing time of converting the SRA read files to the initial experiment Bloom filters. This is a time-consuming process, which took us several days using multiple threads. We did not obtain reliable timing results since we did the conversion on the fly while streaming the data from the SRA over a network connection.

To measure the query time, we first note that there are different use cases that effect how to best measure time performance. SBT approaches are designed to scale to a very large number of experiments or to machines with limited memory (e.g. a desktop computer) because the memory required for a query is not dependent on the number of experiments; only one node of the tree needs to be loaded into memory at any given time. Therefore, we focused our analysis on a setting where the index is loaded into memory with each new query.

Table 2 shows the query speed for all tools. HowDe-SBT was faster than other SBT approaches, with over a 5x speedup on single-transcript batches. Peak RAM usage was < 1.3 GiB for all batches for all tools Table 3.

**Table 2.**
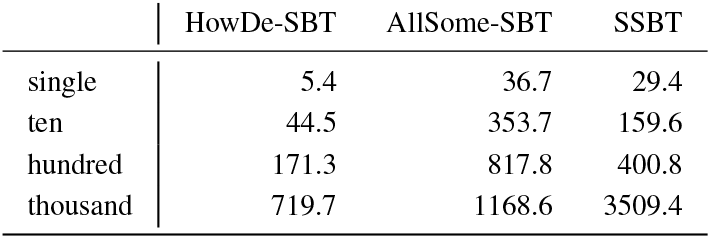
Query times (seconds). Values shown are the median over all the replicates. The cache was cleared prior to each run.

|  | HowDe-SBT | AllSome-SBT | SSBT |
| --- | --- | --- | --- |
| single | 5.4 | 36.7 | 29.4 |
| ten | 44.5 | 353.7 | 159.6 |
| hundred | 171.3 | 817.8 | 400.8 |
| thousand | 719.7 | 1168.6 | 3509.4 |

**Table 3.** Peak resident RAM (in GiB) observed during queries. Values shown are the maximum over all the replicates.

|  | HowDe-SBT | AllSome-SBT | SSBT |
| --- | --- | --- | --- |
| single | 0.8 | 0.3 | 1.3 |
| ten | 0.8 | 0.3 | 1.1 |
| hundred | 0.8 | 0.3 | 1.1 |
| thousand | 0.8 | 0.6 | 1.1 |

We also tested the effect of warming the cache prior to querying, where we ran each query two consecutive times and then reported the run-time for the second run (Supplementary Table S1).

Warming the cache prior to querying led to improved query times for all tools, but their relative performance remained mostly similar.

### 6.3 Bitvector properties

First, we investigate how many positions are active and how saturated the active bits are. The fraction of active *B*_det_ positions decreases going down the tree (by definition), with a median of 0.006 at internal nodes and 0.002 at leaves. The saturation of 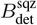 (i.e. the percentage of the active bits that are determined) is 100% at the leaves (by definition) and has a median of 41% at the internal nodes. The saturation of 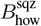 (i.e. the percentage of the active *B*_how_ bits that are 1) has a median of 51% at the leaves and 12% at the internal nodes. It decreases with the height of a node (i.e. maximum distance to a leaf), meaning that at higher levels of the tree, the vast majority of positions that are determined are found to be absent rather than present. Figure 4 shows the saturation distributions. In terms of final space on disk, after RRR compression, the leaves account for only 18% of the total index size.

**Fig. 4.**
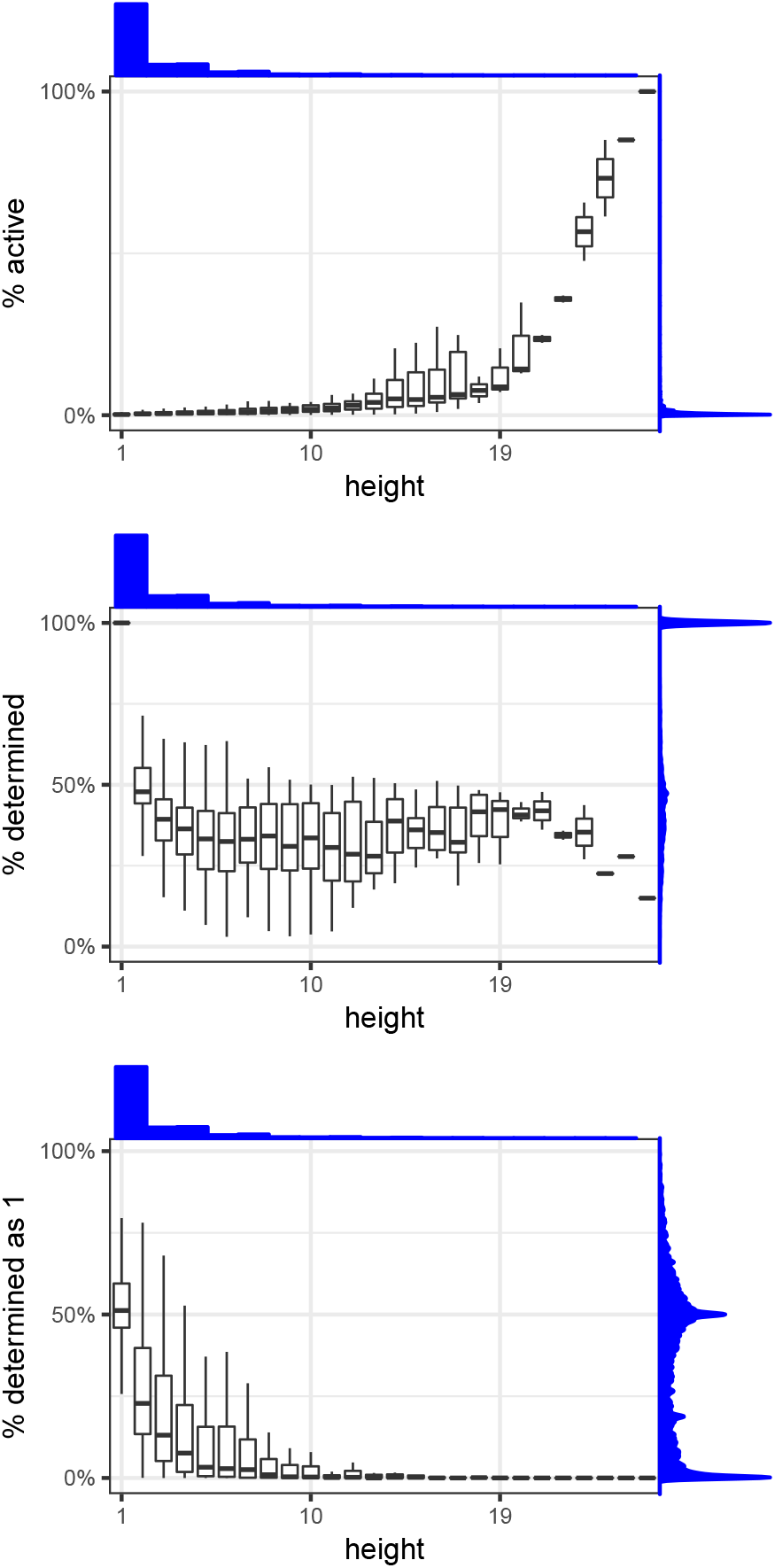
The distribution of nodes’ bitvector saturations, as a function of a node’s height (i.e. maximum distance to a leaf). The top panel shows barplots for the percentages of positions in the *B*_det_ vector that are active, the middle panel shows barplots for the percentages of those positions that are set to 1 (i.e. determined), and the bottom panel shows barplots for the percentages of active *B*_how_ positions that are set to 1. Above each plot is the histogram of height values, and to the right of each plot is the density plot of the percentages being measured in that plot.

In Section 4, we derived the reduction in space and query time for a bit position *x* in terms of the number of nodes where it is active in *B*_det_ (denoted by *n_x_*) and the fraction of nodes where it is set to one in *B*_how_ (denoted by *s_x_*). Figure 3(b) shows the distribution of these values on our tree. The median value for *s_x_* is 0.14 and for *n_x_* is 31, which corresponds to 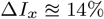 in Figure 3(a). However, we also see a correlation between *n_x_* and *s_x_* (Spearman coefficient of *r* = 0.67). Any future extension of our theoretical analysis per bit position to one of all bit positions jointly should take this complexity into account. We note that the total reduction in index size of HowDe-SBT over SSBT on our data (38%) is due not only to the improved bit representation but also due to the better tree topology that HowDe-SBT constructs; thus, a direct comparison to our theoretical predictions is challenging.

## 7 Conclusion

In this paper, we presented a novel approach for the representation of Sequence Bloom Trees and studied its performance from both a theoretical and an experimental perspective. The main intuition behind our representation is that it reduces the number and entropy of the active bits. Compared to previous SBT approaches, HowDe-SBT is an improvement on all fronts: it constructs the index in less than 36% of the time and with 39% less the space, and can answer small-batch queries five times faster. Compared specifically against AllSome-SBT, the biggest advantage is that the size of the index is an order of magnitude smaller. In comparison against SSBT, the biggest advantage is in the construction time and query times, across all batch sizes.

With the improvements in this paper, the SBT can already be deployed for small and mid-size databases, such as a private patient cohort or all the sequencing data in `flybase.org`. Such a deployment will need to provide an automated way to update the database; while our method naturally supports insertions in 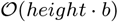 time (we have omitted the details, but Sun *et al.* (2018) give the basic overview of how insertions can be handled), a quality assurance step will be necessary prior to automating database updates. It will also be necessary to provide a front-end interface (e.g. Galaxy or similar to Bradley *et al.* (2017)) for easy access. Such front-ends should also provide wrappers for more biologically-oriented queries (i.e. to convert “which sample has a SNP” into a *k*-mer query).

## Acknowledgements

We are thankful to Ayaan Hossain and Natasha Stopa for helping prototype some aspects of this project. This work has been supported in part by NSF awards DBI-1356529, CCF-551439057, and IIS-1453527 to PM and partially supported by PSU’s College of Engineering Multidisciplinary Seed Grant Program. Research reported in this publication was supported by the National Institute Of General Medical Sciences of the National Institutes of Health under Award Number R01GM130691. The content is solely the responsibility of the authors and does not necessarily represent the official views of the National Institutes of Health.

## Supplementary Section 1 Culling of nodes

In this section, we propose and explore a culling technique for allowing the tree to be non-binary, something that was explored in the original Bloofi paper but was not supported by subsequent SBT approaches. The technique is applied during construction, right after the baseline binary tree topology is constructed but before the bitvector representations are computed.

For a node *u*, let *saturation(u)* be the number of bits set to one and active in *B*_det_(*u*) divided by the number of bits that are active in *B*_det_(*u*). For example, in Figure 1, the left child of the root has saturation of 9/15. Since at this stage Bdet and Bhow are not yet computed, we estimate the saturation by a sub-sampling technique, similar to that used to estimate node similarity First, we identify the *culling threshold* as two standard deviations below the mean of the saturation value in the internal nodes. This was 20% on our dataset. Second, we scan the baseline topology to identify internal nodes *u* for which *saturation*(*u*) is below the threshold. These nodes are then removed from the topology, with their children reassigned as children of the removed node’s parent; if no parent exists, these nodes become roots of a new tree in the forest. We call this process *culling*. The result of culling is that the binary tree potentially becomes a non-binary forest. This does not change the query algorithm in any substantial way.

We investigate the effect of culling on the tree, as a function of the culling threshold (Table S2). At the threshold of 20%, we remove about 4% of the nodes, with a negligible decrease in total index size. The query times fluctuate (Table S3), showing a slight increase or decrease depending on the batch size.

Overall, we conclude that culling did not have a substantial effect on the SBT. It is interesting to observe that when the threshold is high (40%), 45% of the nodes are removed but the index size actually increases by 62%. This is due to the fact that a single active bit in *B*_det_(*u*) is replaced by two active bits in *u*’s children, if *u* is removed.

**Table S1.**
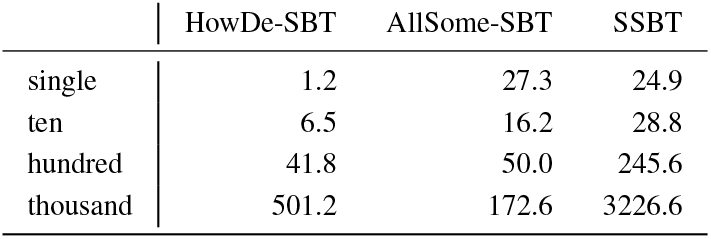
Query times (seconds) using a warm cache. Values shown are the median over all the replicates.

|  | HowDe-SBT | AllSome-SBT | SSBT |
| --- | --- | --- | --- |
| single | 1.2 | 27.3 | 24.9 |
| ten | 6.5 | 16.2 | 28.8 |
| hundred | 41.8 | 50.0 | 245.6 |
| thousand | 501.2 | 172.6 | 3226.6 |

**Table S2.** The effect of culling on the tree.

| Culling threshold | total n. nodes | n. nodes culled | max depth | n. trees in forest | tree size (GiB) |
| --- | --- | --- | --- | --- | --- |
| 0% | 5169 | 0 | 26 | 1 | 14.3 |
| 10% | 5157 | 12 | 25 | 1 | 14.3 |
| 20% | 5056 | 113 | 23 | 2 | 14.3 |
| 30% | 4679 | 490 | 19 | 8 | 14.9 |
| 40% | 3997 | 1172 | 16 | 407 | 23.2 |

**Table S3.** The effect of culling on query time. A cold cache was used.

| Culling threshold | median query time (s) |  |  |  |
| --- | --- | --- | --- | --- |
|  | singles | tens | hundreds | thousands |
| 0% | 5.4 | 45 | 171 | 720 |
| 10% | 5.1 | 43 | 170 | 713 |
| 20% | 6.1 | 39 | 137 | 712 |
| 30% | 7.7 | 44 | 155 | 636 |
| 40% | 7.4 | 44 | 154 | 625 |

## References

Almodaresi, F., Pandey, P., and Patro, R. (2017). Rainbowfish: A succinct colored de Bruijn graph representation. In LIPIcs-Leibniz International Proceedings in Informatics, volume 88. Schloss Dagstuhl-Leibniz-Zentrum fuer Informatik.

Almodaresi, F., Pandey, P., Ferdman, M., Johnson, R., and Patro, R. (2018). An efficient, scalable and exact representation of high-dimensional color information enabled via de Bruijn graph search. bioRxiv, page 464222.

Bloom, B. H. (1970). Space/time trade-offs in hash coding with allowable errors. Communications of the ACM, 13(7), 422–426.

Bradley, P., den Bakker, H., Rocha, E., McVean, G., and Iqbal, Z. (2017). Real-time search of all bacterial and viral genomic data. bioRxiv, page 234955.

Camacho, C., Coulouris, G., Avagyan, V., Ma, N., Papadopoulos, J., Bealer, K., and Madden, T. L. (2009). BLAST+: architecture and applications. BMC bioinformatics, 10(1), 421.

Crainiceanu, A. and Lemire, D. (2015). Bloofi: Multidimensional Bloom filters. Information Systems, 54, 311–324.

Gusfield, D. (1997). Algorithms on strings, trees and sequences: computer science and computational biology. Cambridge University Press.

Holley, G. and Melsted, P. (2019). Bifrost – highly parallel construction and indexing of colored and compacted de bruijn graphs. bioRxiv.

Holley, G., Wittler, R., and Stoye, J. (2015). Bloom filter trie-a data structure for pan-genome storage. In International Workshop on Algorithms in Bioinformatics, pages 217–230. Springer.

Mäkinen, V., Belazzougui, D., Cunial, F., and Tomescu, A. I. (2015). Genome-scale algorithm design. Cambridge University Press.

Marçais, G. and Kingsford, C. (2011). A fast, lock-free approach for efficient parallel counting of occurrences of k-mers. Bioinformatics, 27(6), 764–770.

Muggli, M. D., Bowe, A., Noyes, N. R., Morley, P. S., Belk, K. E., Raymond, R., Gagie, T., Puglisi, S. J., and Boucher, C. (2017). Succinct colored de Bruijn graphs. Bioinformatics, 33(20), 3181–3187.

Mustafa, H., Schilken, I., Karasikov, M., Eickhoff, C., Raetsch, G., and Kahles, A. (2018). Dynamic compression schemes for graph coloring. Bioinformatics, page bty632.

Pandey, P., Almodaresi, F., Bender, M. A., Ferdman, M., Johnson, R., and Patro, R. (2018). Mantis: A fast, small, and exact large-scale sequence-search index. Cell Systems.

Raman, R., Raman, V., and Satti, S. R. (2007). Succinct indexable dictionaries with applications to encoding k-ary trees, prefix sums and multisets. ACM Transactions on Algorithms (TALG), 3(4), 43.

Solomon, B. and Kingsford, C. (2016). Fast search of thousands of short-read sequencing experiments. Nature biotechnology, 34(3), 300–302.

Solomon, B. and Kingsford, C. (2017). Improved search of large transcriptomic sequencing databases using split sequence Bloom trees. In International Conference on Research in Computational Molecular Biology, pages 257–271. Springer.

Sun, C., Harris, R. S., Chikhi, R., and Medvedev, P. (2018). AllSome sequence bloom trees. Journal of Computational Biology, 25(5), 467–479.

Yu, Y., Liu, J., Liu, X., Zhang, Y., Magner, E., Lehnert, E., Qian, C., and Liu, J. (2018). SeqOthello: querying RNA-seq experiments at scale. Genome biology, 19(1), 167.

